## Supplementary figures and images for "E-cadherin mediated Apical Membrane Initiation Site localisation"

### Figure EV1

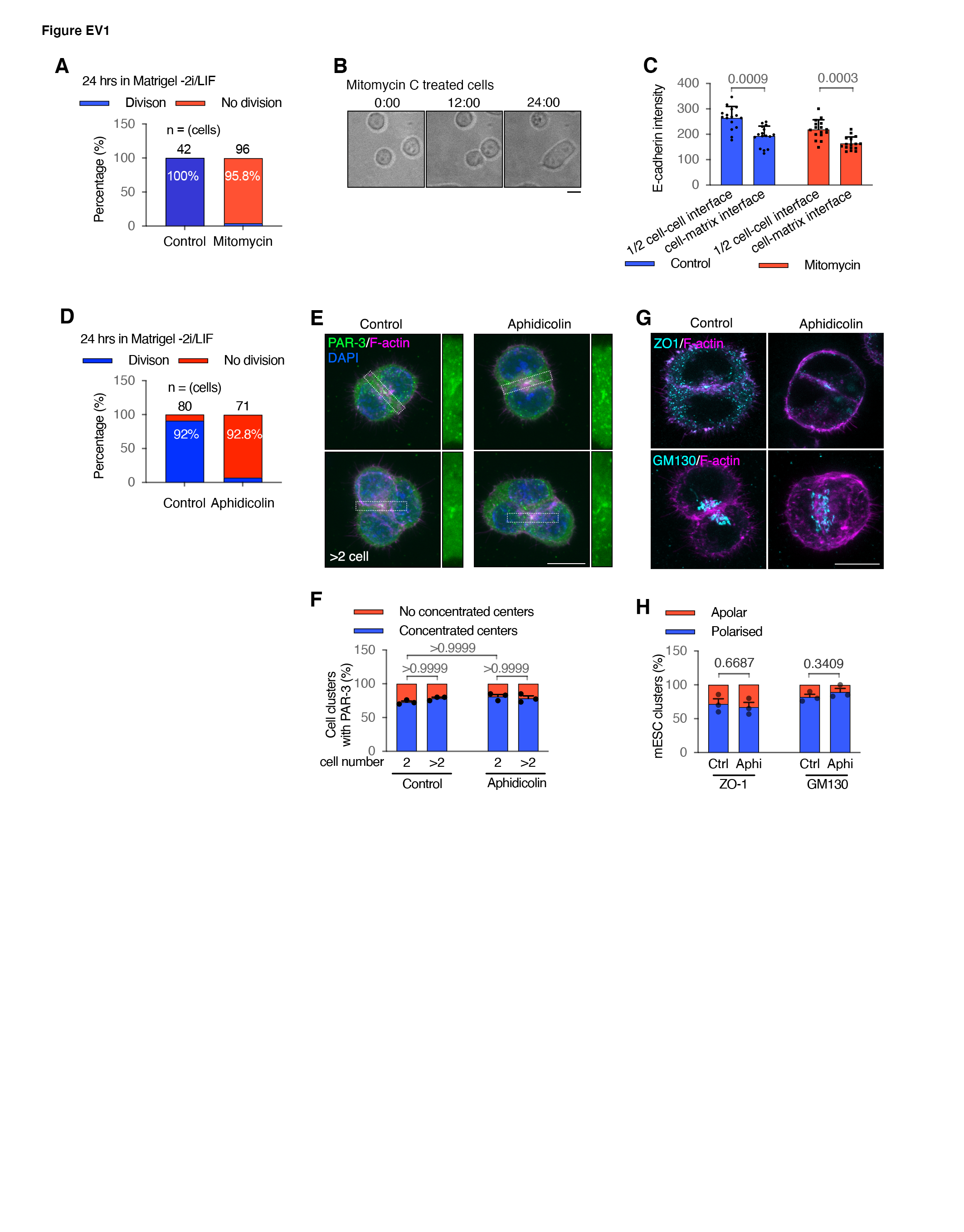

### Figure EV2

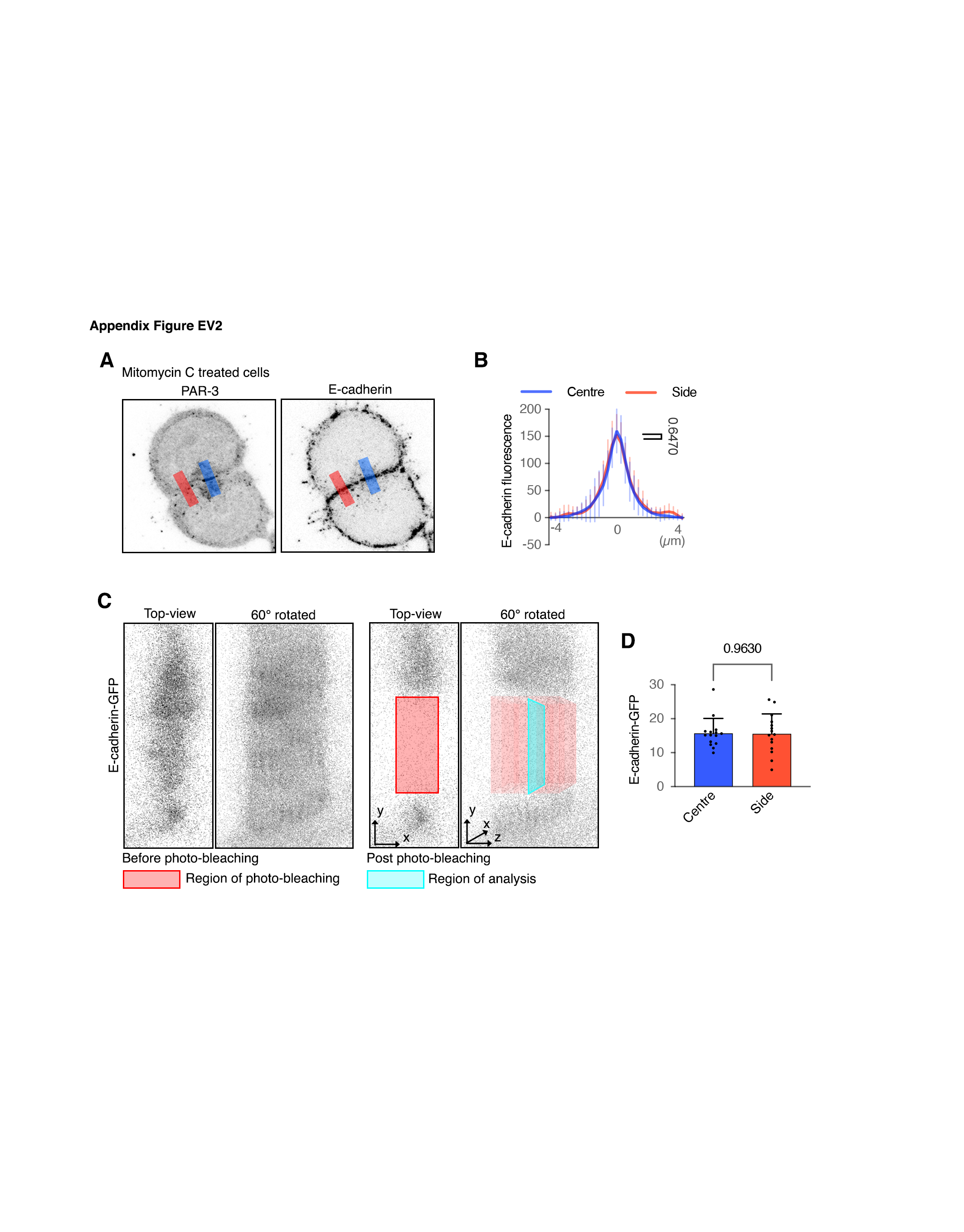

### Figure EV3

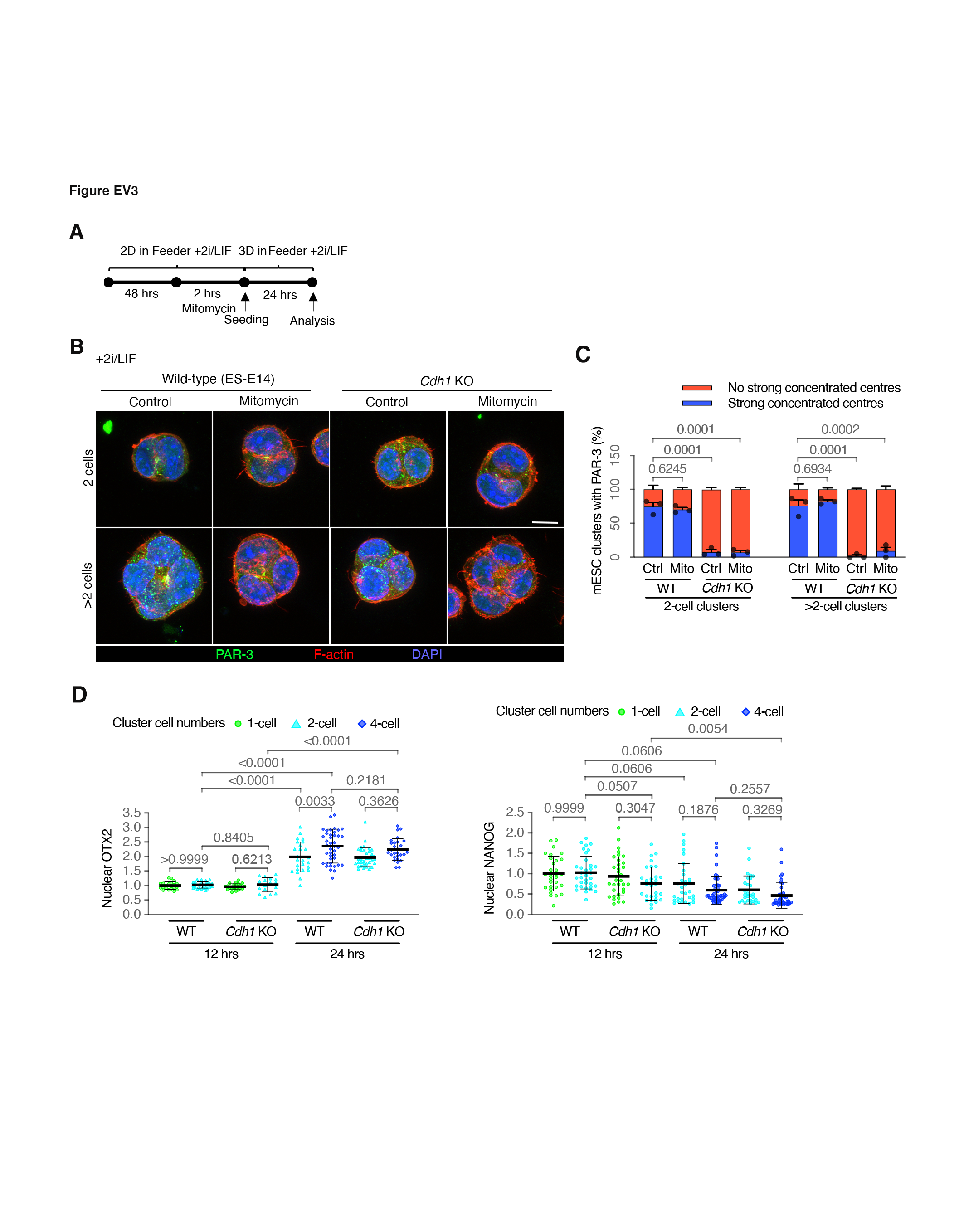

### Figure EV4

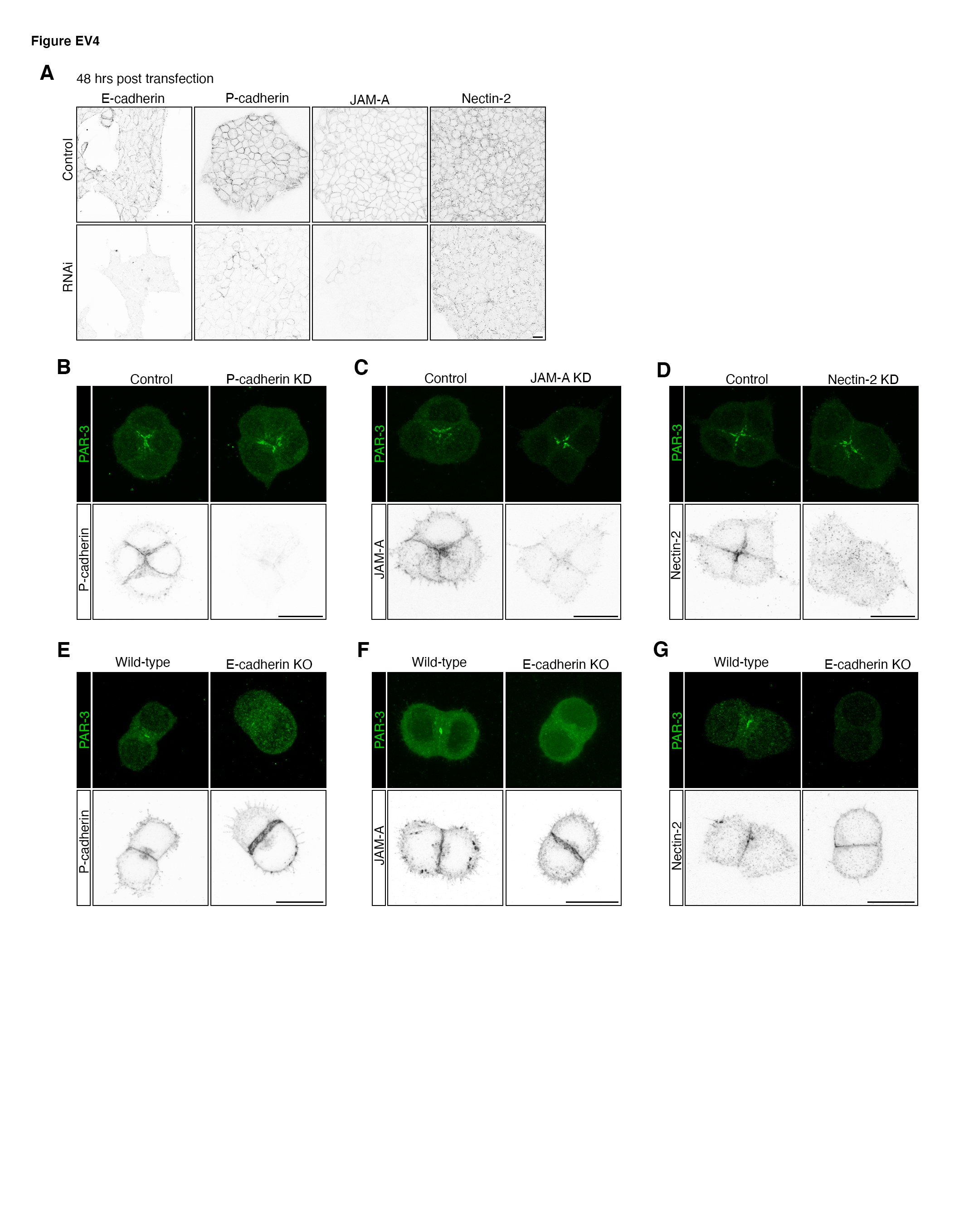

### Figure EV5

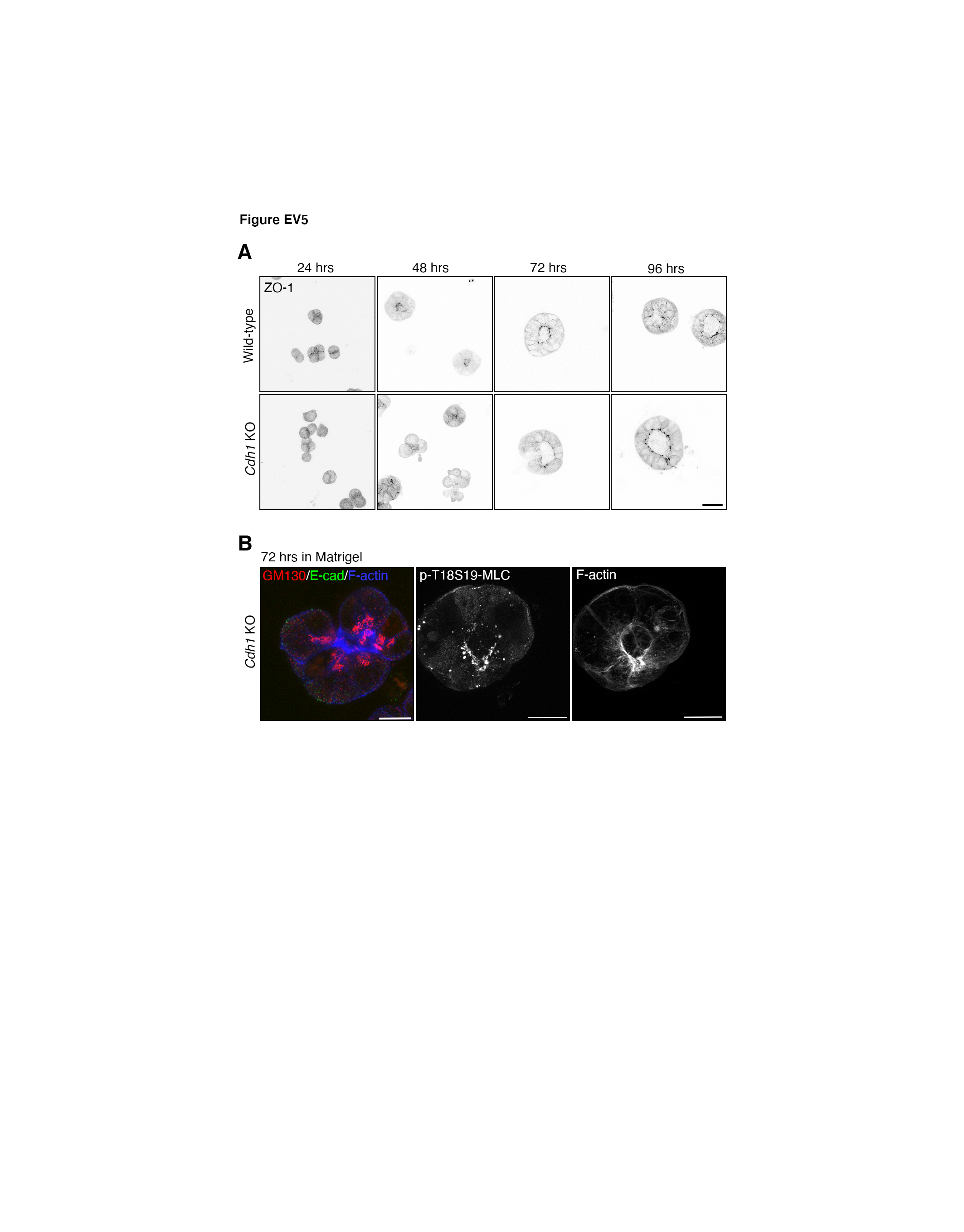
