## Appendix for "E-cadherin mediated Apical Membrane Initiation Site localisation"

**Table of contents:**

- Materials and methods
- Supplementary figures S1-4

### Appendix Material and Methods

#### Aphidicolin treatment

For Aphidicolin treatment, mESCs cultured on gelatin were incubated with 5 μM Aphidicolin (Sc-201535, Insight Biotechnology) in culture media at 37 ℃ for two hours. The treated cells were trypsinised and seeded into 10% Matrigel in culture media containing 5 μM Aphidicolin till further analysis.

#### Line-scan analysis of E-cadherin

To quantify E-cadherin level along the cell membrane in Fig EV1C, the central z-stack from confocal microscope raw images was extracted. On that image, a line of 6 μm wide were drawn perpendicular against the cell-cell membrane or cell-ECM membrane in Fiji. The peak fluorescence value of E-cadherin was extract from the line-scan profile. Three line-scans were done at different regions of the cell-cell membrane in one 2-cell cluster, and three line-scans were done at the cell-ECM membrane in each cell of the 2-cell cluster. The average value of the three or six line-scans was regarded as the E-cadherin level at the cell-cell membrane or the cell-ECM membrane of that 2-cell cluster. To quantify E-cadherin level at the central-most or side regions at the cell-cell interface in Fig EV3B, a line of 3 μm wide was drawn perpendicular to the cell-cell interface to generate the line-scan profile. The region corresponding to the polarised PAR-3 dot was determined as the central-most region, and regions between the PAR-3 centre and cell-free regions was determined as the side region. Two side regions were analysed at each side flanking the centre region, and the average line-scan profile of the two was taken as the line-scan profile at the side region of a 2-cell cluster.

#### Analysis of nuclear OTX2 or NANOG

To quantify nuclear OTX2 or NANOG level, raw images were captured on the confocal microscope with OTX2 or NANOG and DAPI channels. The raw images that contained z-axis stacks were subjected to maximum value projections in Fiji. The DAPI channel was turned into a binary image and the region under DAPI signals were masked by using the Analyse Particles tool in Fiji. The total OTX2 or NANOG fluorescence values in each cell that were under the correspondent DAPI mask was measured and regarded as the OTX2 or NANOG level of that cell. Based on cell cluster morphologies displayed by F-actin signals and nucleus morphologies displayed by DAPI signals, only cells at the interphase were analysed.

### Appendix Supplementary Figures

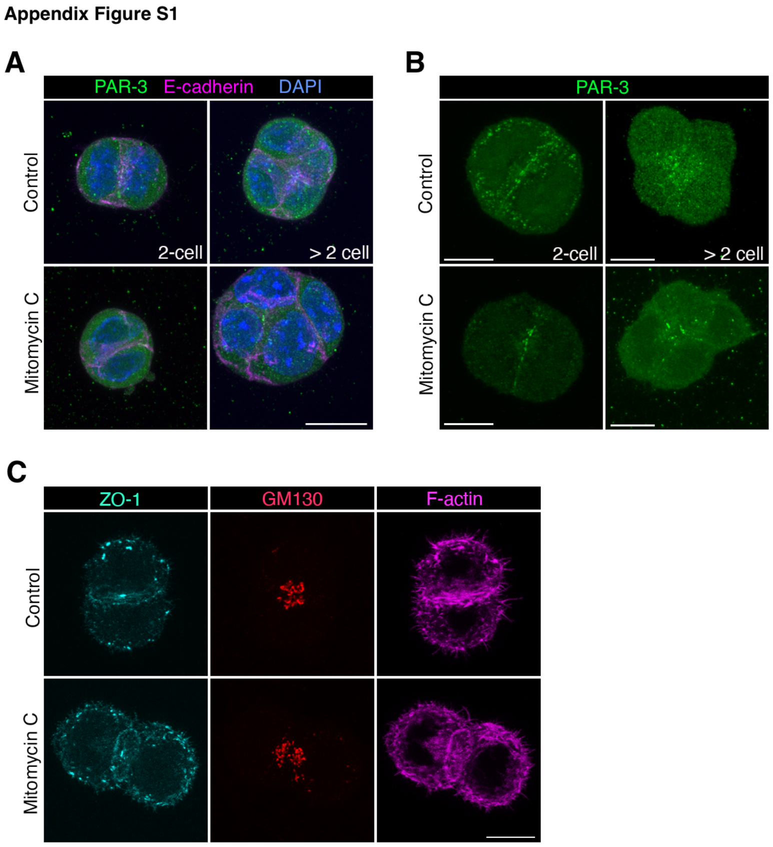


**Appendix Fig S1 - Immunofluorescence of PAR3, ZO-1 and GM130 mouse embryonic stem cells related to Figure 1.**

A Representative images of mESCs that did not localise PAR-3 to their centres.

B mESC clusters that had weak or non centralised PAR-3 along cell-cell interfaces.

C Split-channel images of Figure 1H. All scale bars: 10 µm.

Appendix Figure S2

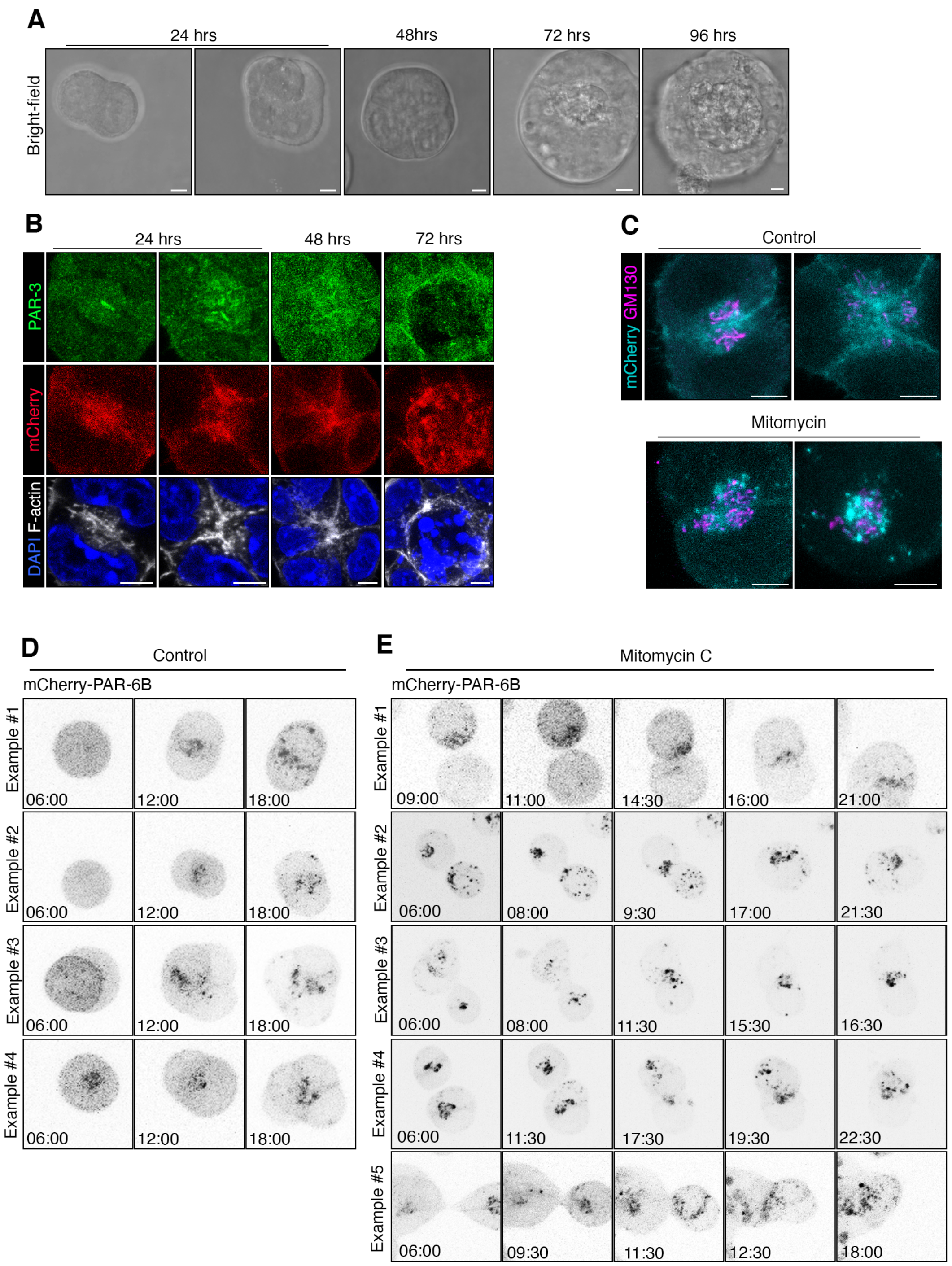


**Appendix Fig S2 - Polarisation and lumenogenesis in the mCherry-PAR6B mouse embryonic stem cell line.**

A Bright-field images of Fig 2A.

B Immunofluorescence of PAR-3 in the mCherry-PAR6B mouse embryonic stem (mESC) line cultured 24-72 hours in Matrigel.

C Immunofluorescence of GM130 in the mCherry-PAR6B mESC line cultured 24 hours in the Matrigel. mCherry is partially co-localised with the Golgi network but is largely located at sub-apical or apical membrane regions. All scale bars: 5 µm.

D,E Example images from movies of mCherry-PAR6B control (D) and mitomycin treated (E) mESCs cultured in Matrigel.

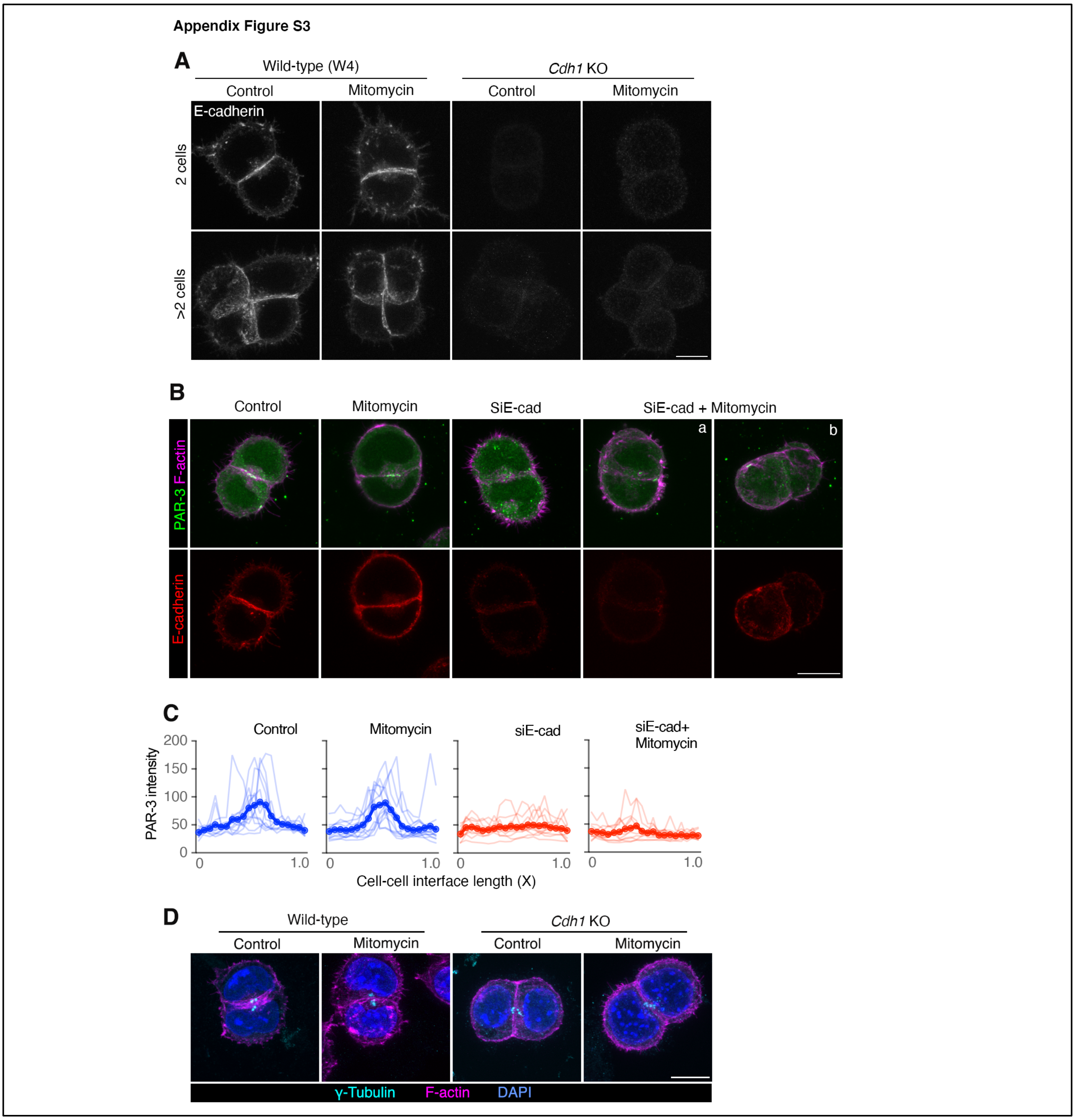


**Appendix Fig S3 - Wild-type, E-cadherin knock-down and knock-out cell clusters.**

A Images related to Figure 3A. Immunofluorescence of E-cadherin in wild-type (W4) and E-cadherin knock-out (*Cdh1* KO) mESCs.

B, C Immunofluorescence of PAR-3 and E-cadherin (B) and line-scans of PAR-3 at the cell-cell interface of 2-cell mESC doublets (C) in wild-type (ES-E14) and E-cadherin siRNA knock-down mESCs cultured for 24 hours in Matrigel, with and without cell division. Line-scans were sectioned and fitted to each 5% along the cell-cell interface length. N = 12-15 lines scans for each condition.

D Representative images of centrosomes in wild-type and *Cdh1* KO mESCs cultured 24 hours in Matrigel with or without cell divisions.

All scale bars: 10 µm.

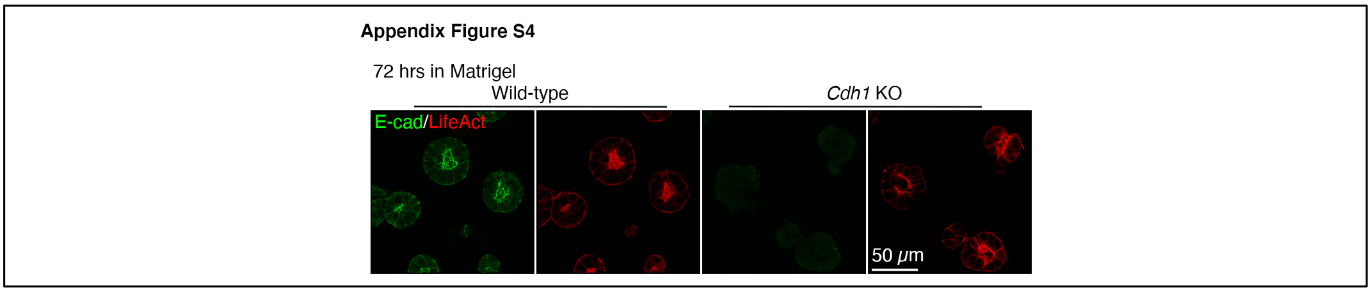


**Appendix Fig S4. Immunofluorescence of E-cadherin in the LifeAct-mRuby WT and *Cdh1* KO mESC lines cultured for 72 hours in Matrigel.**
